# Hi-C informed kernel association test: integrating 3-dimensional genome structure into variant-set association for whole-genome sequencing data

**DOI:** 10.1101/2025.10.28.684891

**Authors:** Yueyang Huang, Riddhik Basu, Wenbin Lu, Shannon T. Holloway, Yun Li, Jung-Ying Tzeng

**Affiliations:** Bioinformatics Research Center, North Carolina State University, Raleigh, NC, USA; Department of Statistics, North Carolina State University, Raleigh, NC, USA; Department of Population Health Sciences, Duke University, Durham, NC, USA; Department of Biostatistics, University of North Carolina at Chapel Hill, Chapel Hill, NC, USA; Department of Genetics, University of North Carolina at Chapel Hill, Chapel Hill, NC, USA

**Author notes:** Corresponding author: Jung-Ying Tzeng, Department of Statistics and Bioinformatics Research Center, North Carolina State University, Raleigh, NC, USA.

## Abstract

Variant-set association analysis is a powerful strategy for genetic studies of whole genome sequence (WGS) data, especially for rare variants. By aggregating variant signals, variant-set analysis can improve statistical power, result interpretability, and study replicability. Motivated by evidence that three-dimensional (3D) genome architecture plays a critical role in regulating gene transcription, several works have incorporated 3D genome architecture into gene-based association tests and demonstrated great promise. In this work, we extend the idea of 3D-genome guided test from gene-centric to gene-agnostic, whole-genome testing by introducing a Hi-C informed kernel association test. We present a principled procedure that converts Hi-C contact confidence into borrowing weights and integrates these weights into genetic similarity kernels so that higher-confidence interacting loci contribute more to the association test of the target variant set. We use a controlling parameter to adaptively determine the appropriate degree of information borrowing from its interacting loci during association testing. We assess the performance of the Hi-C informed test using simulations and illustrate its advantage in detecting rare-variant sets using WGS data from the ARIC study in the Trans-Omics for Precision Medicine (TOPMed) program.

## Introduction

Whole-genome sequencing (WGS) enables comprehensive analysis of all types of genetic variants across the entire genome, with particular strength in detecting rare variants. In human genome, rare variants constitute the majority of genetic variants [1] and are known to contribute to the etiology of many complex diseases [2, 3, 4]. For rare variant association studies, variant-set analysis provides an attractive alternative to single-variant analysis due to its ability to jointly evaluate the effects of multiple variants within a set (e.g., gene) and increase power to detect association signals in aggregate. Rare-variant tests can be broadly classified into two classes: burden tests, which assume variants influence the trait in the same direction and with similar strength [5, 6, 7], and variance component (kernel-based) tests, which model variant effects as random and can accommodate heterogeneous directions and magnitudes of effects across variants [8, 9, 10, 11, 12, 13].

Recent studies have shown that incorporating biological and functional information of variants can further enhance the performance of variant-set analyses [e.g., 14, 15]

Besides functional annotation, integrating three-dimensional (3D) genome architecture has also been shown to improve the power and interpretability of variant-set analyses [16, 17, 18]. The 3D genome architecture plays a critical role in gene regulation [19, 20]. In the eukaryotic nucleus, chromosomal DNA is folded into highly condensed and organized 3D structures, allowing transcriptional and regulatory processes to function efficiently within distinct domains [21, 22]. The 3D folding brings regions that are distant on the linear genome into close spatial proximity, enabling direct physical contact between regions. Consequently, non-coding regulatory elements (e.g., enhancers and silencers) can modulate gene transcription by acting on gene promoters located far away in linear distance. A well-known example is the obesity-associated locus within an intron of *FTO*, which physically interacts with and regulates *IRX3* through enhancer activity [23, 24, 25, 26]. Variants within *IRX3* also associate with obesity [27, 28, 29], supporting the notion that risk variants in close 3D spatial proximity may jointly influence complex traits. Finally, disruption of 3D genome organization has also been linked to various diseases [30, 31], which further highlights the functional importance of 3D genome structure in human complex traits.

A number of chromosome conformation capture assays have been developed to investigate the 3D genome architecture [19, 32, 33, 20]. In particular, highthroughput chromosome conformation capture (Hi-C) adopts a genome-wide approach and captures all chromatin contacts between DNA regions in the nucleus simultaneously. Hi-C begins with formaldehyde crosslinking, which preserves interactions by chemically bonding two DNA molecules in close spatial proximity. The cross-linked DNA is then digested with a restriction enzyme, and the resulting DNA fragments are ligated to form a chimeric DNA that represents physical contacts. These chimeric DNA fragments are subsequently sheared and ligated to sequencing adapters to create a Hi-C library. The library is then sequenced using high-throughput sequencing technologies to identify interacting DNA regions.

Focusing on gene-based association test, H-MAGMA [18] uses Hi-C chromatin contact data to link distal regulatory elements to gene promoters, and aggregates association signals from all SNPs assigned to the gene, including SNPs located within the gene body, promoter, and linked distal regulatory regions. [17] develops a powerful multi-component gene-based testing framework that incorporates long-range chromatin interaction, functional annotations, and flexible definitions of genic and regulatory region sizes. In this framework, the gene-based variant sets are defined by incorporating putative regulatory elements identified from ChIP-seq data and the activity-by-contact (ABC) model. The resulting variant sets used for gene-level association tests include SNPs located in the gene body as well as proximal and distal SNPs residing in putative regulatory elements. Both H-MAGMA and the framework of [17] demonstrate the utilities of integrating 3D chromatin architecture to improve gene-level association tests.

In this work, we extend these ideas by introducing a Hi-C informed kernel association test that adaptively integrates 3D genomic information into variant-set association analysis. We develop the Hi-C informed test under the kernel machine (KM) regression framework because it includes both burden test and SKAT test [12] as special cases and can be naturally extended to the omnibus version, SKAT-O [34]. Our method differs from previous approaches in two key aspects. First, instead of restricting analyses to gene regions and focusing on the corresponding long-range interactions among gene promoters and regulatory elements, our approach considers gene-agnostic, whole-genome variant-set analysis across the entire genome, where we partition the genome into variant sets aligned with Hi-C data and incorporate SNP information from loci that significantly interact with the target variant set. Second, when evaluating the association of the target variant set, information from its interacting loci is incorporated according to the statistical confidence (e.g., q-value) of the corresponding Hi-C contacts. The contribution from interacting loci is adaptively weighted by the trait of interest, ensuring that only 3D interactions relevant to the trait under study are included. We evaluate the performance of our Hi-C informed test through extensive simulation studies using real WGS data from the Trans-Omics for Precision Medicine (TOPMed) Program of of the National Heart, Lung, and Blood Institute. Finally, we apply the Hi-C informed method to WGS data from the Atherosclerosis Risk in Communities (ARIC) study and identify promising genomic regions associated with platelet counts.

## Methodology

### Hi-C Informed Kernel Association Test

As Hi-C contact data are typically available at 10-kb resolution, we define each variant set as the variants located within a 10-kb genomic region. Following the Hi-C literature [35], we also refer to each 10-kb genomic region as a “locus”. As illustrated in Figure 1, our workflow consists of four main steps: **(1)** Obtain locus–locus contact information in the form of q-values from Hi-C experiments; **(2)** convert contact q-values to locus-locus weight values, which determine the degree of information borrowing from the neighboring loci that significantly interact with the target locus; **(3)** construct the Hi-C informed kernel function by incorporating the locus–locus weights so that variants in loci with stronger contact signals contribute more to genetic kernel similarity; and **(4)** conduct the Hi-C informed kernel association test, which evaluates the association of the target locus with the trait while adaptively determining the optimal borrowing degree from its interacting loci. Below we describe each step in detail.

**Figure 1.**
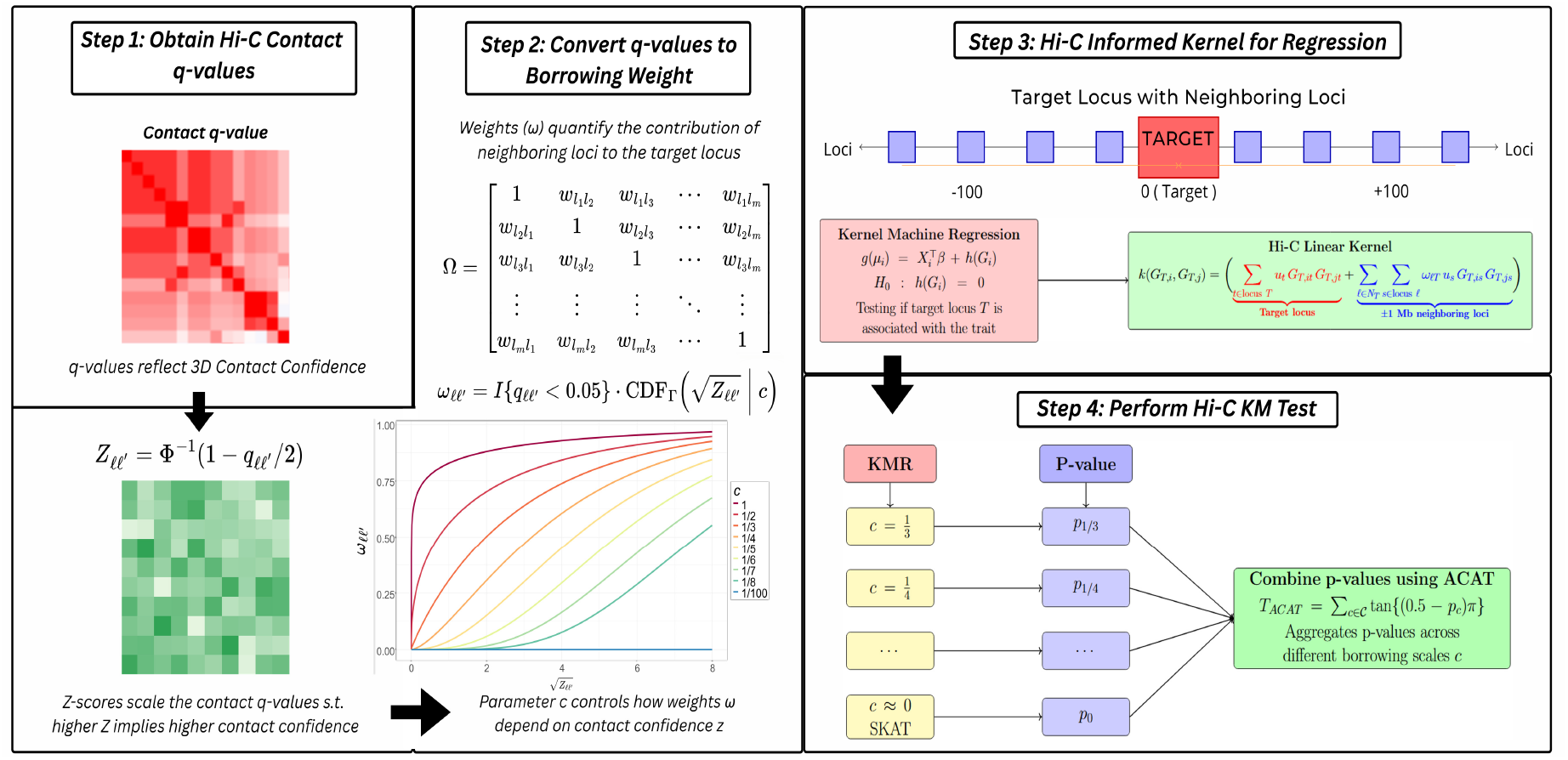
Overview of the Hi-C Kernel Association Test

#### Step 1. Obtain the Hi-C Locus-Locus Contact Information (q-values)

We begin by retrieving Hi-C data downloaded from HUGin [36]. Instead of using raw contact counts between locus pairs, we construct the Hi-C contact matrix using the corresponding q-values, which quantify the statistical significance of spatial proximity between locus pairs. The q-values are computed by Fit-Hi-C [37], a method that effectively distinguishes biologically meaningful chromatin contacts from those resulting from random polymer looping. Fit-Hi-C assigns q-values to chromatin contacts based on Hi-C sequencing results, adjusting for genomic distance and technical biases via spline regression. These q-values reflect the statistical confidence that two genomic loci are in close 3D spatial proximity, and hence provide a more functionally relevant measure of chromatin interaction than raw contact counts.

#### Step 2. Convert the Locus-Locus Contact q-values to Locus-Locus Weight Values

Given a q-value between two loci *𝓁* and *𝓁*^′^, denoted as *q*_*𝓁𝓁*_*′*, we convert it to a weight value *ω*_*𝓁𝓁*_*′* ∈ [0, 1], which quantifies the degree to which information is borrowed from locus *𝓁′* when assessing the association at locus *𝓁*. This transformation is conducted in two step. First, we transform the q-value into a Z-score using: ^*Z*^*𝓁𝓁*^′^ = Φ^−1^(1 − *q 𝓁𝓁*^′^ *′*/2), where Φ^−1^ is the inverse cumulative distribution function (CDF) of the standard normal distribution. We divide q-value by 2 so that *q*_*𝓁𝓁*_*′* /2 < 0.5, which ensures the transformed Z score to be positive. Second, for a locus pair with significant Hi-C interaction (i.e., *q*_*𝓁𝓁*_*′ <* 0.05), we map the Z score to a weight value *ω*_*𝓁𝓁*_*′* ∈ [0, 1], where 1 indicates full borrowing of information from locus *𝓁*^′^.

As shown in Table 1, the distribution of *Z*_*𝓁𝓁*_*′* ‘s for locus pairs with q-values *<*0.05 is highly right-skewed, ranging from 1.96 to 38.47 with a median of 4.27. To reduce the influence of extreme values and compress the range, we take a square root and then use the Gamma CDF to map the compressed Z-value into *ω*_*𝓁𝓁*_*′* within the [0, 1] interval:

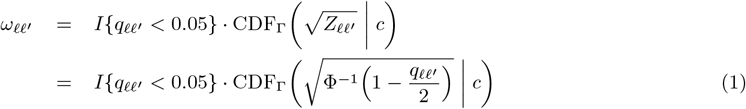

where CDF_Γ_(· | *c*) is the CDF of Gamma distribution with mean 1*/c*. We use the Gamma CDF because of its positive support and flexibility in modeling skewed distributions. Compared to alternatives such as the standard normal CDF or logistic function, the Gamma CDF provides greater control over the transformation behavior. Specifically, by tuning the shape and scale parameters of the Gamma distribution (or equivalently, its mean and variance), we can regulate how rapidly the borrowing weight *ω*_*𝓁𝓁*_*′* increases with 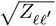 and how quickly *ω*_*𝓁𝓁*_*′* approaches 1. This allows for a more flexible and adaptive borrowing scheme based on the spatial proximity (as captured by q-values) between loci.

**Table 1:** Five-number summary of contact confidence *Z*_*𝓁𝓁*_*′* = Φ^−1^(1 − *q*_*𝓁𝓁*_*′*/2) and 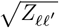 for locus pairs *𝓁* and *𝓁′* with Hi-C contact q-value *q*_*𝓁𝓁*_*′ <* 0.05.

|  | Min. | 1st<br>Quantile | Median | 3rd<br>Quantile | Max. |
| --- | --- | --- | --- | --- | --- |
| $Z_{\ell\ell'}$ | 1.96 | 2.92 | 4.27 | 6.70 | 38.47 |
| $\sqrt{Z_{\ell\ell'}}$ | 1.40 | 1.71 | 2.07 | 2.59 | 6.19 |

To regulate how much information locus *𝓁* borrows from other loci, we use a controlling parameter *c*, defined such that the Gamma distribution has mean 1*/c* and fixed variance 10. As illustrated in Figure 2, a large *c* value (e.g., *c* = 1) promotes broader borrowing, i.e., allowing loci with weaker spatial contacts (i.e., smaller *Z*_*𝓁𝓁*_*′*) to still contribute to the association test at locus *𝓁*. In contrast, a smaller *c* value (e.g., *c* = 1/8) leads to more localized borrowing, i.e., only from loci with stronger contact evidence (i.e., larger *Z*_*𝓁𝓁*_*′*), and the maximum borrowing weight is also reduced. Although lim*c*→0 *ω*_*𝓁𝓁*_*′* = 0, the Gamma distribution is undefined at *c* = 0. Therefore in our implementation, we use a small value (e.g., *c* = 1/100) to obtain *ω*_*𝓁𝓁*_*′* = 0 ∀ *𝓁*^′^ ≠ *𝓁*, and the test reduces to the original SKAT [12], which evaluates each locus without incorporating any information from spatial interacting loci.

**Figure 2.**
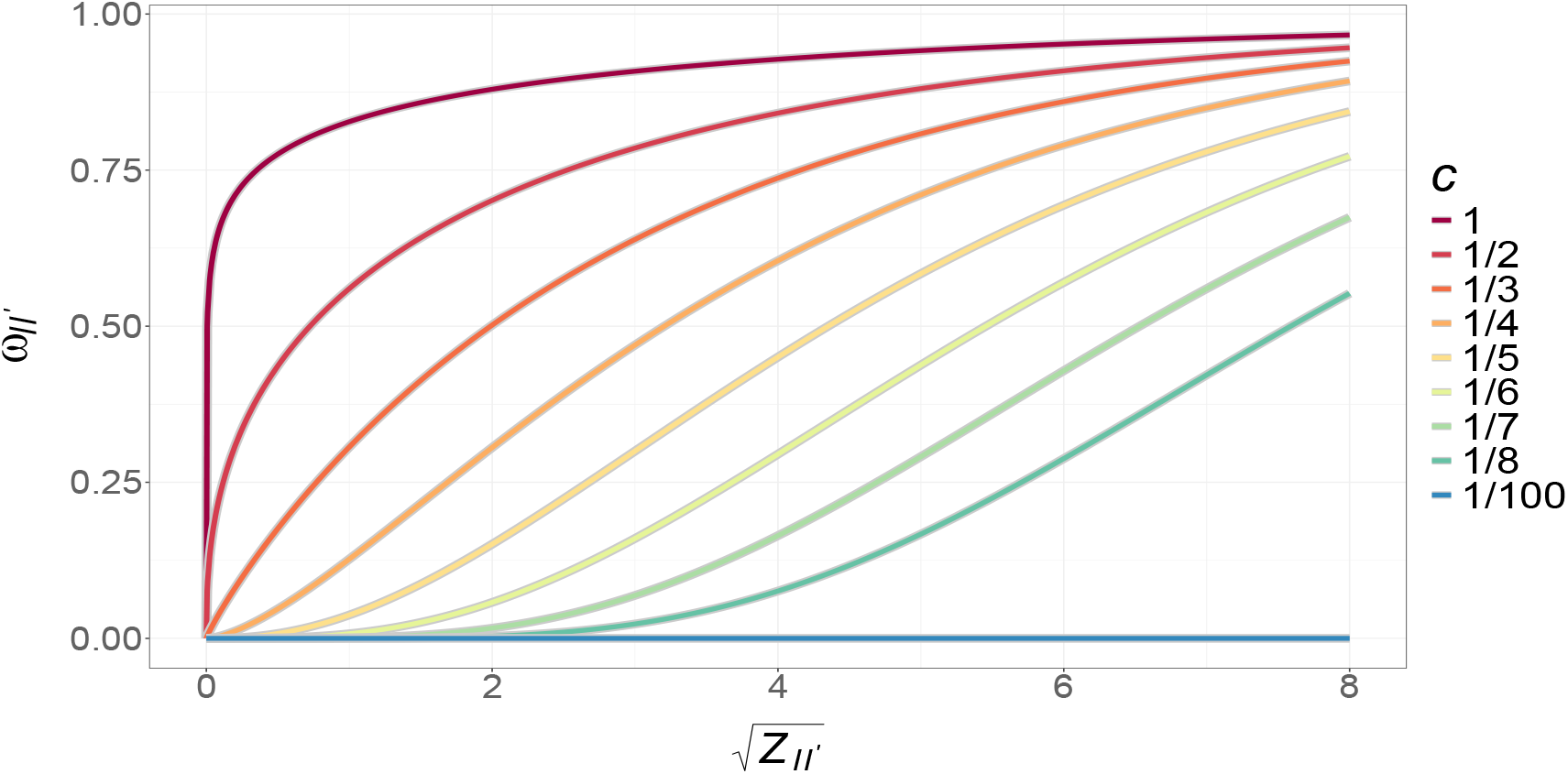
Relationship between locus contact confidence 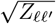 and borrowing weights *ω*_*𝓁𝓁*_*′* under varying *c* values, showing how *ω*_*𝓁𝓁*_*′* changes with 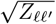 *′* for different values of the parameter *c*. The transformation is based on a Gamma CDF with mean 1*/c* and fixed variance 10.

We make two additional remarks. First, after extensive empirical exploration, we choose the parameterization with mean 1*/c* and variance 10, as it yields a desirable *Z*-*ω* relationship for moderating the degree of borrowing based on contact q-values. While other parameter configurations are possible, we found this configuration to be empirically effective in our numerical analyses based on real WGS data from TOPMed. Second, in practice, we consider a grid of *c* values and let the data determine the most informative borrowing scheme (i.e., the *c* value minimizing the test p-value) [38, 39]. This approach provides additional adaptivity and allows the Hi-C spatial information to be incorporated in a data-driven manner into the association test.

#### Step 3. Construct the Hi-C Informed Kernel for KM Regression

Given a “target locus” *T* (i.e., the 10-kb region to evaluate association with the trait), we use a KM regression framework to assess the association between the trait and the variants within the target locus *T*, after “optimally” incorporating information from its interacting loci. KM regression captures complex, potentially nonlinear relationships by implicitly mapping the data into a higher-dimensional feature space, where the association can be represented by a linear model.

In Hi-C informed KM test, we consider the ±1Mb region (i.e., ± 100 loci) of the target locus *T* as the “neighboring loci” and incorporate the contact structure between the target locus and its neighboring loci into the association test. We focus on the ±1Mb neighboring window because regulatory elements such as enhancers can influence trait-associated genes over distances up to 1Mb upstream or downstream of their target [40, 41, 42] Furthermore, the Hi-C data also show that more than half of the significant genomic interactions (q-value *<*0.05) are within 1Mb distance, as seen in Figure 3.

**Figure 3.**
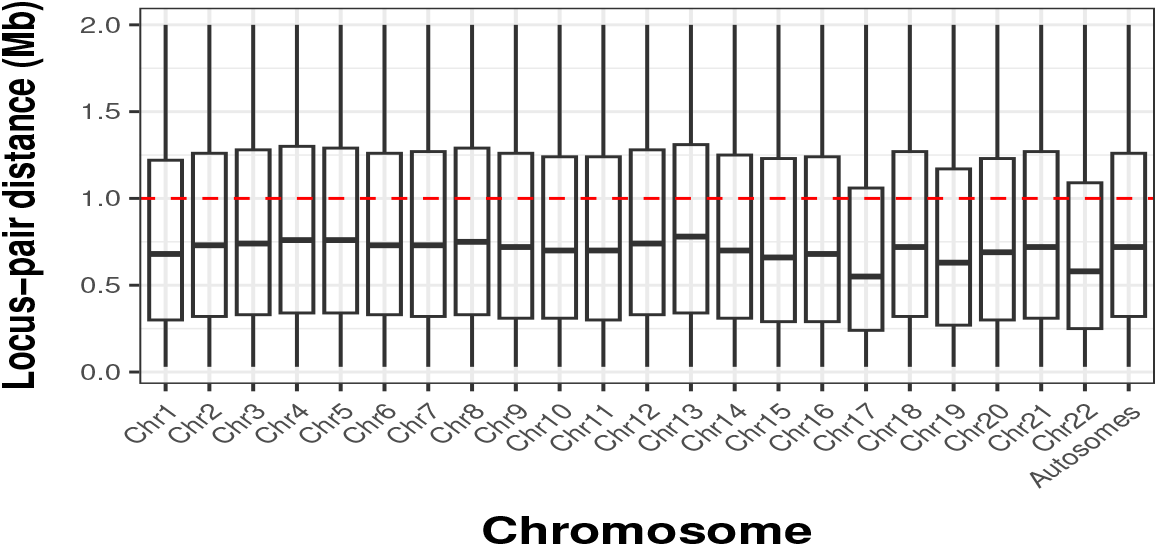
Boxplots of distance between locus pairs with significant Hi-C interactions (i.e., q-value*<* 0.05). Distances are measured between the midpoints of two loci.

Therefore, for a target locus *T*, we obtain the contact q-values of its neighboring loci and convert them to the contact weight values *ω*_*𝓁𝓁*_*′* using Equation (1). The resulting weights for all 100 + 1 + 100 = 201 loci are stored in the locus-locus weight matrix **Ω**_*T*_ ∈ ℝ^201×201^ for later use.

Assume that the target locus contains *M*_*T*_ variants and that the left and right neighboring windows contain *M*_*L*_ and *M*_*R*_ variants, respectively. The total number of variants for KM test of target locus *T* is *M* = *M*_*T*_ + *M*_*L*_ + *M*_*R*_. Then for subject *i*, let *G*_*T,i*_ ∈ ℝ^*M*^ be the genotype vector of the *M* variants in the target locus and its Hi-C neighbors, *y*_*i*_ ∈ ℝ^1^ be the trait value, and *X*_*i*_ ∈ ℝ^*p*^ be the covariate vector. Then the KM regression is given as

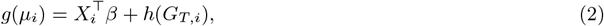

where *g*(·) is a known link function (e.g., the identity link for continuous traits or the logit link for binary traits); *µ*_*i*_ = *E*(*y*_*i*_ | *X*_*i*_, *G*_*T,i*_) is the expected trait value given *X*_*i*_ and *G*_*T,i*_; *β* is the *p ×* 1 coefficient vector for the covariates; and *h*(*G*_*T,i*_) is a smooth function representing the genetic effect. By the representer theorem [43], *h*(*G*_*T,i*_) can be written as 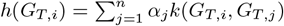, where *k*(·, ·) is a kernel function measuring genetic similarity between subjects *i* and *j*, and *α*_*j*_ ‘s are unknown coefficients. Equivalently, in matrix form,

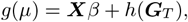

where *µ* = [*µ*_1_, · · ·, *µ*_*n*_]^⊤^ ∈ ℝ^*n*^, ***X*** = [*X*_1_, · · ·, *X*_*n*_]^⊤^ ∈ ℝ^*n×p*^, ***G***_*T*_ = [*G*_*T*,1_, · · ·, *G*_*T,n*_]^⊤^ ∈ ℝ^*n×M*^, and *h*(***G***) = ***K***_*T*_ *α*. Here ***K***_*T*_ = {*k*_*ij*_ } is the *n × n* kernel matrix with entries *k*_*ij*_ = *k*(*G*_*T,i*_, *G*_*T,j*_), and *α* = [*α*_1_, · · ·, *α*_*n*_]^⊤^. The kernel machine model, Equation (2), has been shown to be equivalent to a random effects model:

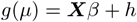

with *h* ∼ *N* (0, *τ*_*T*_ ***K***_*T*_) [10, 9]. Therefore, testing for the association between trait and genetic variants in locus *T*, i.e., *H*_0_ : *h*(***G***_*T*_) = 0, is equivalent to testing *H*_0_ : *τ*_*T*_ = 0.

The key step of Hi-C informed KM test lies in the construction of the kernel function *k*(*G*_*T,i*_, *G*_*T,j*_). The underlying rationale is that, when computing genetic similarity between subjects *i* and *j*, we assign non-trivial weights only to variants in loci that significantly interact with the target locus, with higher weights assigned to loci exhibiting stronger interaction signals. This can be done by incorporating the locus-level weights *ω*_*𝓁𝓁*_*′* stored in **Ω**_*T*_ into the kernel function as described below. The computational complexity arises primarily from the need to map the locus-level weights into corresponding weights at the variant level. To illustrate, consider the original burden kernel function as

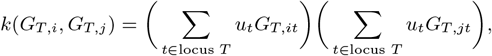

where *G*_*T,it*_ is the genotype of variant *t* for subject *i*, and *u*_*t*_ is the variant-specific weight (e.g., based on minor allele frequencies (MAF) such that rarer variants receive greater weights). The burden kernel is the product the weighted genotype sum for subject *i* and that for subject *j*. The Hi-C informed burden kernel extends the original burden kernel by incorporating all variants within the ± 1Mb neighboring loci into the kernel calculation, and weighting the neighboring variants according to their interaction with the target locus *T* as quantified in the locus-locus weights **Ω**_*T*_. Specifically, we index the neighboring locus by *𝓁*, with *𝓁* ∈ {−100, −99,· · ·, −1} for the 100 loci in the left neighboring window and *𝓁* ∈ {1, 2, · · ·, 100} for the 100 loci in the right neighboring window. The Hi-C informed burden kernel is then

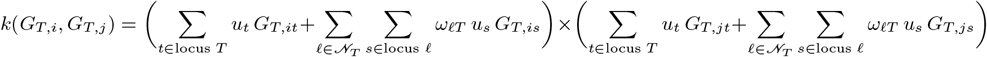

where 𝒩_*T*_ = {−100,· · ·, −1, 1, · · ·, 100} is the index set of the neighboring locus of the target locus *T* ; note that locus *T* itself is excluded from the neighboring set 𝒩_*T*_ ; and *ω*_*𝓁T*_ is the (*𝓁, T*)-entry of the locus-locus weight matrix **Ω**_*T*_.

Similarly, the original linear kernel is

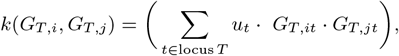

which is the weighted sum of the genotype products between subjects *i* and *j* over all loci. Unlike the burden kernel, which assumes homogeneous effects across variants, the linear kernel can accommodate heterogeneous variant effects and provide greater flexibility and robustness in modeling genetic architecture. The Hi-C informed linear kernel function becomes

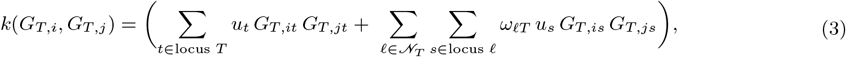

where variants from neighboring loci are incorporated, with interaction-based weights *ω*_*𝓁T*_ determining their contribution to the overall genetic similarity.

#### Step 4. Perform Hi-C Informed Kernel Association Test

The association between the trait and the target locus *T* can be assessed by testing the null hypothesis *H*_0_ : *τ*_*T*_ = 0. We use the score-like test [12]

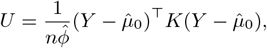

where 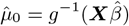 is the fitted trait mean under *H*_0_, 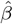 is the estimated covariate coefficient under *H*_0_, and 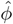 is the dispersion parameter estimate under *H*_0_. For continuous trait, *ϕ* is the residual variance under *H*_0_ and for binary trait, *ϕ* = 1.

For a fixed *c, U* follows a weighted 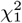 distribution asymptotically under *H*_0_ [12, 44, 45] under *H*_0_, and the corresponding p-value, denoted by *p*_*c*_, can be computed using the Davies method [46] or the moment-matching approach [47]. Since the optimal borrowing scale *c* is not known, we consider a grid of *c* ∈ 𝒞 ≡ {*c*_1_, *c*_2_, · · ·, *c*_*max*_}, compute the p-value for each *c* in grid 𝒞, and then combine the *p*_*c*_’s using the aggregated Cauchy association test (ACAT) [48]:

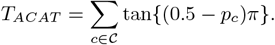

The ACAT statistic, *T*_*ACAT*_, behaves similarly to the minimum p-value test statistic because *T*_*ACAT*_ is dominated by the smaller *p*_*c*_’s. However, unlike the minimum p-value methods, the ACAT p-value can be analytically approximated using the Cauchy distribution, even when the input p-values are correlated as in our case:

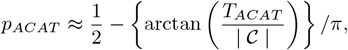

where | 𝒞 | is the number of element in grid 𝒞.

### Simulation Design

We conduct simulation studies with two main objectives: (i) to evaluate the performance of the proposed Hi-C informed KM test in detecting association signal of the target locus, and (ii) to determine an appropriate specification of the grid 𝒞 and its maximum value *c*_*max*_. We use the WGS data of the ARIC study, which was sequenced under the Trans-Omics for Precision Medicine (TOPMed) program.

We partition each chromosome into 10 kb regions (hereafter referred to as ‘loci’) to align with the Hi-C data. For each locus, we collect the Hi-C information for its 200 neighboring loci within the ±1 Mb window, Including 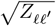 and the number of interacting loci, i.e., the number of loci with which the locus has significant 3D interactions (i.e., *q*_*𝓁𝓁*_*′ <* 0.05).

For simulation studies, we focus on data from chromosome 5 (chr5), which was chosen because its distributions of 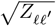 values and the number of interacting loci are similar to those observed in the whole-genome data (Table 2). Based on the information, we randomly select three loci as “target loci”, which have 30, 60, and 100 interacting loci, corresponding to the first, second, and third quantiles of the empirical distribution of the number of interacting loci per locus (Table 2). Given a target locus, we randomly select 15 of its interacting loci, and then randomly designate 10% of the variants within the target locus and each of these 15 loci as causal.

**Table 2:** Comparisons of Hi-C features between whole-genome and chromosome 5 data based on (A) the number of interacting loci per locus, (B) 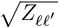 *′* within a chromosome, and (C) 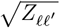 within ±1 Mb. Loci *𝓁* and *𝓁′* are considered as interacting if their Hi-C contact q-value *q*_*𝓁𝓁*_*′ <* 0.05. 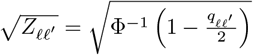.

|  | Minimum | 1st Quartile | Median | 3rd Quartile | Maximum |
| --- | --- | --- | --- | --- | --- |
| **(A) Number of interacting loci per locus** |  |  |  |  |  |
| Whole Genome | 1 | 27 | 64 | 100 | 185 |
| Chromosome 5 | 1 | 30 | 68 | 104 | 180 |
| **(B) $\sqrt{Z_{\ell\ell'}}$ for locus pairs within the same chromosome** |  |  |  |  |  |
| Whole Genome | 1.401 | 1.709 | 2.066 | 2.588 | 6.185 |
| Chromosome 5 | 1.401 | 1.710 | 2.068 | 2.589 | 6.185 |
| **(C) $\sqrt{Z_{\ell\ell'}}$ for locus pairs within $\pm 1$ Mb** |  |  |  |  |  |
| Region selected | 1.401 | 1.783 | 2.279 | 2.851 | 6.185 |
| Chromosome 5 | 1.401 | 1.818 | 2.260 | 2.862 | 6.185 |

In addition, we simulate two covariate variables, a continuous covariate *X*_1*i*_ generated from *N* (0, 1) and a binary covariate *X*_2*i*_ generated from Bernoulli(0.5). Using the covariates as well as the causal variants from the target locus and its 15 selected interacting loci, we simulate continuous trait values *Y*_*i*_ using the following model:

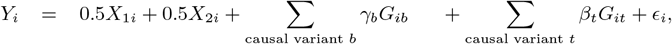

where *ϵ*_*i*_ ∼ *N* (0, 1); *G*_*ib*_ is the genotype of causal variant *b* in the 15 interacting loci with effect size *γ*_*b*_; and *G*_*it*_ is the genotype of causal variant *t* in the target locus with effect size *β*_*t*_.

#### Type I Error Simulations

We consider two settings for Type I error simulations: Setting I to evaluate whether the proposed Hi-C informed test preserves the Type I error rate at the nominal level 0.05 when neither the target nor its interacting loci have causal effects, and Setting II to evaluate the magnitude of Type I error inflation when the interacting loci contain causal variants but the target locus does not. In Setting I, we set both *γ*_*b*_ and *β*_*t*_ to zero, so that no genetic variants influence the trait. The Setting I model reduces to

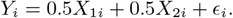

In Setting II, we set *β*_*t*_ = 0 and simulate *Y*_*i*_ using the model

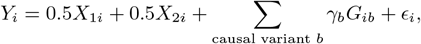

where the effect size of each neighboring causal variant is defined as

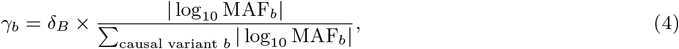

MAF_*b*_| with MAF_*b*_ denoting the MAF of variant *b* and δ_*B*_ controlling the overall effect size of the interacting causal variants. Equation (4) is designed such that rarer causal variants have larger effects while the overall background effect remains fixed at δ_*B*_. We examine three levels of *“background effects”* from the interacting loci in the ±1 Mb neighborhood, corresponding to δ_*B*_ ∈ {11, 12.5, 14}, for small, medium, and large effects, respectively.

#### Power Simulations

To evaluate power, we set both *β*_*t*_ ≠ 0 and *γ*_*b*_ ≠0 in Equation (4) when gener ating *Y*_*i*_. For causal variants in the target locus, the effect sizes, *β*_*t*_’s, are set as *β*_*t*_ = 12.5 × | log_10_ MAF_*t*_|∑ _causal variant *t*_ | log_10_ MAF_*t*_|. For the neighboring causal variants, we use the same *γ*_*b*_ as in Setting II of the Type I error simulation, i.e., δ_*B*_ ∈ {11, 12.5, 14}, corresponding to background effects that are small (i.e., smaller than target effects), medium (i.e., same as target effects), and large (i.e., larger than target effects), respectively.

Table 3 summarizes the simulation designs. We use the original SKAT (i.e., *c*_*max*_ = 1/100) as the baseline method to evaluate the performance of the Hi-C informed test in detecting the association at the target locus. We implement the Hi-C informed test using the linear kernel (Equation (3)) with the MAF weights specified by the beta density as in SKAT [12], i.e.,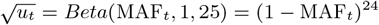. We implement the Hi-C informed test by considering different *c*_*max*_ in the grid of *c*, i.e., 𝒞 = {1/100, 1/8, 1/7, 1/6, 1/5, 1/4, 1/3, · · ·, *c*_*max*_}. Specifically, we evaluate *c*_*max*_ = 1/3, 1/4, 1/5, 1/6, 1/7, and 1/8. In all simulation scenarios, we perform 1000 replicates and evaluate the performance at *α* = 0.05 using sample sizes of *n* = 6, 000 and *n* = 9, 000.

**Table 3:** Overview of simulation designs and purposes. All simulations are performed with 1,000 replicates using sample sizes of *n* = 6, 000 or 9, 000.

| Simulation | Causal Predictors | Goal |
| --- | --- | --- |
| Type I Error - Setting I | Covariates only | Assess false positive rates when no genetic effects are present |
| Type I Error - Setting II | Covariates<br>+ Interacting loci* | Assess false positive rates when interacting loci in the $\pm 1$ Mb neighboring region have effects but the target locus does not |
| Power | Covariates<br>+ Interacting loci*<br>+ Target locus** | Assess power to detect target locus when both target and interacting loci have effects |

\* : Three levels of background effect sizes from the interacting loci within the $\pm 1$ Mb neighboring region are considered: small, intermediate, and large.
\*\* : Three levels of interactivity are considered for the target locus: 30, 60, and 100 interacting loci within the $\pm 1$ Mb neighboring region of a target locus.

## Results

### Simulation Results

#### Type I Error Simulations - Setting I

Table 4 shows the empirical Type I error rates of the Hi-C informed test in Setting I (no effects from the target locus and its interacting loci) for *c*_*max*_ = 1/3, 1/4, 1/5, 1/6, 1/7, and 1/8. The results suggest that the Type I error rate is controlled at the nominal level of 0.05 across different *c*_*max*_ values, sample sizes, and numbers of interacting loci of the target locus, although with a slightly conservative trends similar to SKAT.

**Table 4:**
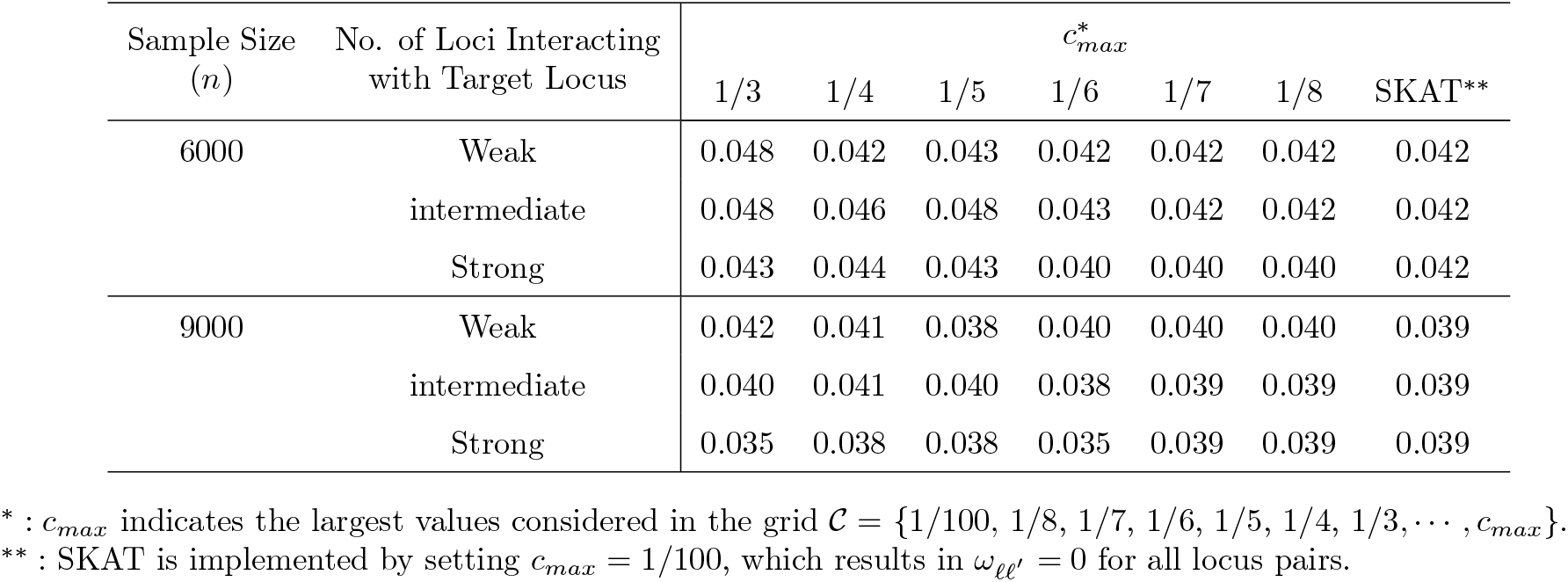
Type I error rates in Simulation Setting I (no causal effects from the target locus and its interacting loci) at the nominal level of 0.05.

| Sample Size<br>($n$) | No. of Loci Interacting<br>with Target Locus | $c_{max}^*$ | | | | | | |
| --- | --- | --- | --- | --- | --- | --- | --- | --- |
|  |  | 1/3 | 1/4 | 1/5 | 1/6 | 1/7 | 1/8 | SKAT** |
| 6000 | Weak | 0.048 | 0.042 | 0.043 | 0.042 | 0.042 | 0.042 | 0.042 |
|  | intermediate | 0.048 | 0.046 | 0.048 | 0.043 | 0.042 | 0.042 | 0.042 |
|  | Strong | 0.043 | 0.044 | 0.043 | 0.040 | 0.040 | 0.040 | 0.042 |
| 9000 | Weak | 0.042 | 0.041 | 0.038 | 0.040 | 0.040 | 0.040 | 0.039 |
|  | intermediate | 0.040 | 0.041 | 0.040 | 0.038 | 0.039 | 0.039 | 0.039 |
|  | Strong | 0.035 | 0.038 | 0.038 | 0.035 | 0.039 | 0.039 | 0.039 |

\* : $c_{max}$ indicates the largest values considered in the grid $\mathcal{C} = \{1/100, 1/8, 1/7, 1/6, 1/5, 1/4, 1/3, \dots, c_{max}\}$ .
\*\* : SKAT is implemented by setting $c_{max} = 1/100$ , which results in $\omega_{\ell\ell'} = 0$ for all locus pairs.

#### Type I Error Simulations - Setting II

Setting II (no effects from the target locus but causal effects from its interacting loci) allows us to evaluate the extent of Type I error inflation of the Hi-C informed test under different numbers of interacting loci of the target locus (30, 60, and 100) and different magnitudes of background effects from the interacting loci (small, intermediate, and large). The findings, together with the power performance, can inform the appropriate range for *c*_*max*_.

The results are shown in Figure 4 for *n* = 6, 000 and Figure 5 for *n* = 9, 000. When *n* = 6, 000 and the target locus has 30 interacting loci (top row of Figure 4), we observe that *c*_*max*_ = 1/3 and 1/4 always yield inflated Type I error rates, and the inflation becomes more pronounced as the background effect sizes from the neighboring loci increase (i.e., from left to right). Similar inflation patterns are observed when the number of interacting loci increases to 60 (middle row) and 100 (bottom row), except that *c*_*max*_ = 1/5 begins to show moderate inflation, although the inflation is less severe than at *c*_*max*_ = 1/3 and 1/4. Comparable results are also observed for *n* = 9, 000 in Figure 5, although the inflation at *c*_*max*_ = 1/5 is more pronounce. Overall, these results show that Hi-C informed KM test can preserve Type I error rate across a broad range of conditions when *c*_*max*_ ≤ 1/6, and often also when *c*_*max*_ = 1/5.

**Figure 4.**
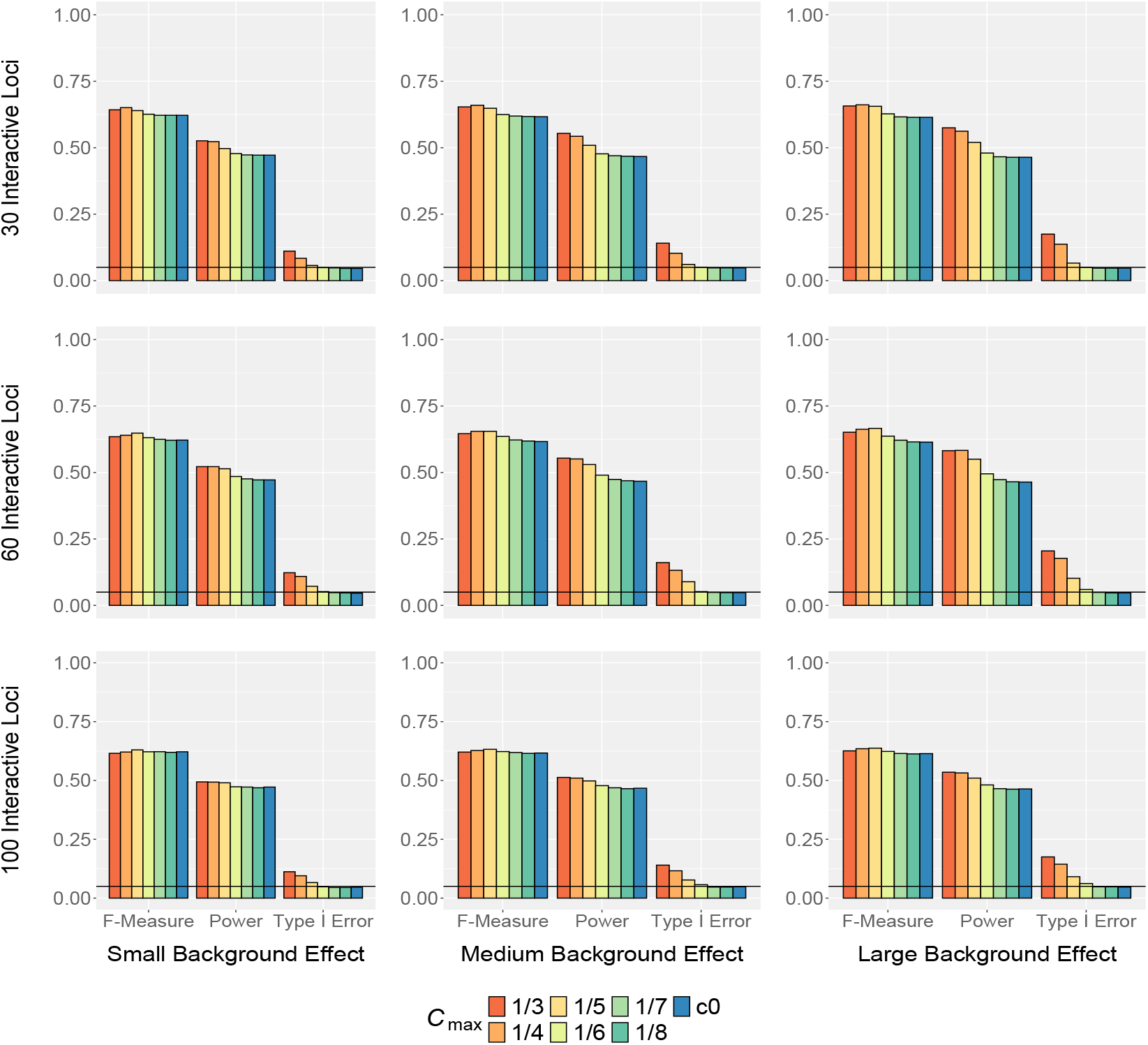
Simulation results for sample size *n* = 6, 000 showing type I error rate, power, and f-measure. Type I error rate is calculated when the target locus has no causal effect but its interacting loci do (i.e., Setting II of the type I error Simulation). Power is calculated when both the causal locus and its interacting loci have causal effects.

**Figure 5.**
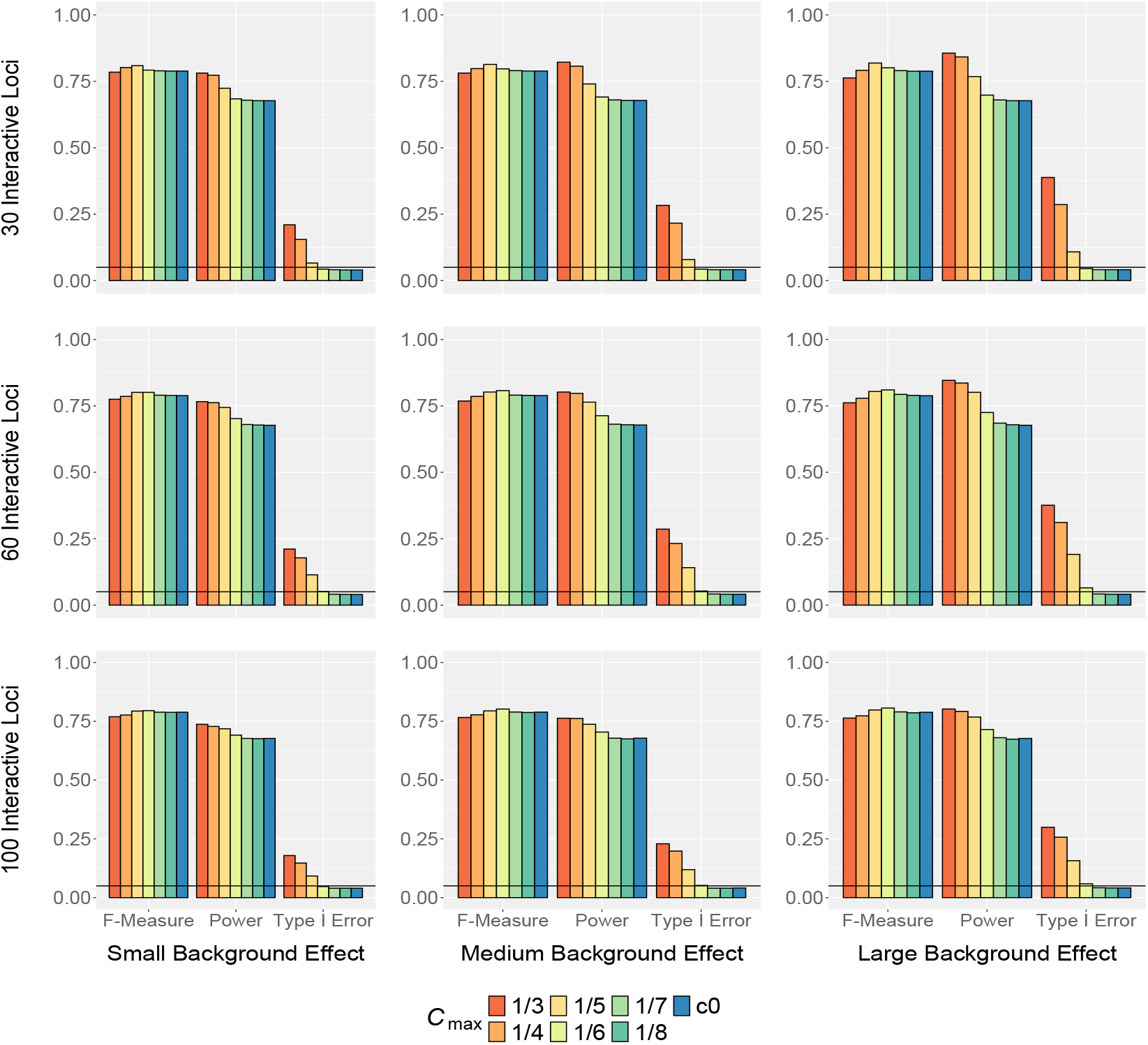
Simulation results for sample size *n* = 9, 000 showing type I error rate, power, and f-measure. Type I error rate is calculated when the target locus has no causal effect but its interacting loci do (i.e., Setting II of the type I error Simulation). Power is calculated when both the causal locus and its interacting loci have causal effects.

#### Power Simulations

We observe a general trend that power decreases as *c*_*max*_ becomes smaller. The trend holds across different numbers of interacting loci and different magnitudes of background effects from neighboring loci, suggesting the power gain from incorporating information from interacting loci. However, larger *c*_*max*_ does not necessarily yield higher power. For example, both *c*_*max*_ = 1/3 and *c*_*max*_ = 1/4 often have similarly high power regardless of the sample size, but the Type I error rates for *c*_*max*_ = 1/3 are higher than those for *c*_*max*_ = 1/4. These observations suggest that incorporating excessive neighboring information may dilute the association signals of the target locus, resulting in little or no additional power gain. At the same time, it can also increase the risk of false positives. Therefore, careful selection of *c*_*max*_ is essential to balance power gain with Type I error control. On the other hand, some *c* grid values may be too small to effectively leverage neighboring information. For example, the power of *c*_*max*_ = 1/8 is nearly identical to SKAT, both yielding the lowest power. This suggests that 1/8 could be excluded from the grid 𝒞.

#### Identifying Suitable *c*_*max*_ via Power-Type I Error Tradeoffs

Because power gains may arise at the cost of inflated Type I error rates, we also evaluate a composite measure (referred to as “f-measure”) following the spirit of the classical F-measure. The f-measure is defined as

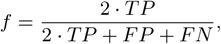

where *TP* and *FN* are the numbers of rejections and non-rejections, respectively, out of 1,000 repetitions in the power simulations; and *FP* is the number of rejections out of 1,000 repetitions in Setting II of the Type I error simulation. We observe that the highest f-measure occurs at *c*_*max*_ = 1/5 in most scenarios with *n* = 6, 000 and in some scenarios with *n* = 9, 000. In other cases with *n* = 9, 000, *c*_*max*_ = 1/6 achieves the highest f-measure. These results suggest that *c*_*max*_ = 1/5 or 1/6 provides an optimal trade-off between power gains from incorporating information of interactive loci and the risk of inflated false positives. Based on the assessment of Type I error rate, power, f-measure across various scenarios, we suggest using the grid 𝒞 = {1/100, 1/7, 1/6, 1/5} in practice.

### Application to the ARIC Study

We apply the Hi-C informed test to identify rare variant sets (i.e., MAF*<* 0.01) that are associated with platelet count in individuals of European ancestry using the WGS data from the TOPMed ARIC study. The ARIC study is a prospective cohort study designed to investigate the etiology of atherosclerotic diseases. DNA samples were sequenced at approximately 30× coverage, with genotype calls based on the TOPMed Freeze 8 release. We remove SNPs with Hardy-Weinberg Equilibrium (HWE) p-values *<* 10^−14^ after adjusting for population structure, and exclude one member of each individual pair with kinship coefficient *>* 0.177 (i.e., first-degree relations). There are 6,260 individuals of European ancestry with both WGS genotypes and platelet counts for downstream analysis.

We covert the TOPMed WGS data from GRCh38 to GRCh37 using UCSC liftOver to align with the Hi-C data coordinates. We then partition the genome into 10 kb regions, each treated as a locus, and use the locus midpoint as its representative coordinate to align with the Hi-C contact map. For each locus, we perform variant-set analysis using the Hi-C informed test with grid 𝒞 = {1/100, 1/7, 1/6, 1/5} and the original SKAT (implemented with *c* = 1/100). Both tests use the linear kernel and variant-specific weight of Beta(MAF, 1, 25), i.e., 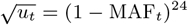. In all analyses, we adjust for age, age^2^, sex, and the first 10 principal components (PCs) to account for population substructure. Because the distribution of platelet counts is skewed, we apply rank-based inverse normal transformation as did in [48]. There are 233,349 loci across the autosome, and hence we set the genome-wide significance threshold at 2.14 × 10^−7^ = 0.05/233, 349 using the Bonferroni method.

Table 5 shows the loci identified as significant by at least one of the two methods. Both the Hi-C informed test (p-value 8.33 × 10^−8^) and original SKAT test (p-value 5.72 × 10^−8^) identify a significant locus located in 16,200-16,210 kb on chromosome 19, overlapping the *TPM4* gene. *TPM4* has been previously implicated in platelet count and volume, and insufficient expression of *TPM4* in megakaryocytes is known to impair platelet production [49]. The slightly larger p-value of the Hi-C informed test compared with SKAT shows the cost of evaluating a grid of *c* values when the association signal can already be identified by original SKAT.

**Table 5:** Significant Loci in the ARIC analysis.

| Chromosome | Region (kb) | Gene | Hi-C Test p-value | SKAT p-value |
| --- | --- | --- | --- | --- |
| 19 | 16,200–16,210 | *TPM4* | $8.33 \times 10^{-8}$ | $5.72 \times 10^{-8}$ |
| 9 | 4,830–4,840 | *RCL1* | $1.36 \times 10^{-7}$ | $8.36 \times 10^{-7}$ |

Hi-C informed test additionally identifies another significant locus located in 4,830-4,840 kb on chromosome 9 (p-value 1.36 × 10^−7^) within the *RCL1* gene. Common variants in *RCL1*, e.g., rs13300663, have previously been reported to be associated with platelet traits [50, 51, 52, 53]. Within the grid of 𝒞, *c* = 1/6 yields the smallest p-value. At this setting of *c* = 1/6, three loci have their *ω*_*t𝓁*_ *>* 0.2 (as listed in Table 6) among the 81 significant chromatin contact loci (q-value *<* 0.05) within the ±1 Mb window of the target locus. All three loci are located within the *JAK2* gene. Previous studies have reported a gene fusion between *JAK2* and *RCL1* in cervical squamous cell carcinoma [54]. Elevated platelet counts have been identified as a prognostic factor in cervical cancer [55], suggesting a potential mechanistic link. Although chromatin contacts suggest potential 3D interactions between *RCL1* and *JAK2*, the association analysis finds significance at *RCL1* but not *JAK2*. This pattern might indicate that the signal arises primarily from *RCL1*, with *JAK2* potentially involved through long-range interactions.

**Table 6:**
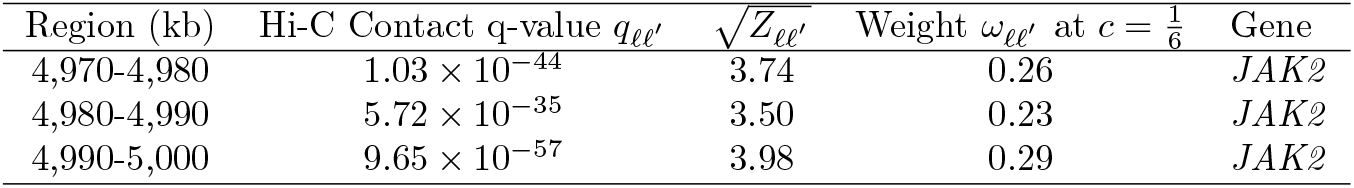
Loci with substantial interactions (i.e., *ω*_*𝓁𝓁*_*′ >* 0.2) with Locus 9:4830–4840 in *RCL1* in the ARIC analysis.

## Discussion

In this work, we present a framework to incorporates 3D genome architecture into variant-set association tests. We illustrate strategies to translate data from high-throughput chromosome conformation capture (Hi-C) assays into locus-specific weights. These weights reflect the confidence of 3D interactions between the target locus and its neighboring loci within a ±1 Mb region, and specify how much information each interacting locus contributes to the association test of the target locus. In this framework, only loci with significant Hi-C interactions are considered, and the final weights are modulated by a tuning parameter *c* that is determined by the data. Information from interacting loci is borrowed only when supported by the data. Through numerical studies using both simulations and real data application, we show potential gains in power and improvements in overall performance accounting for false positives and false negatives, as quantified by the f-measure.

Our work provides a proof of concept for a whole-genome testing strategy that extends beyond gene regions to leverage 3D chromatin interaction information. Our implementation focuses on continuous traits, applies the SKAT framework with a linear kernel, and adopts a Gamma CDF based weighting approach to incorporate Hi-C data. While these specifications demonstrate the feasibility and utility of the Hi-C informed test, future work is needed to extend to other trait types, different testing approaches, and alternative strategies for integrating 3D genome information.

Another limitation of the current study is that locus resolution of the association test is restricted to 10 kb segments. As Hi-C technology advances and higher resolution data become available through deeper sequencing, the test resolution can be further improved. The proposed framework remains applicable regardless of the resolution.

Finally, the performance of Hi-C informed test may be affected by the accuracy and reliability of Hi-C data. Hi-C contact maps could be noisy and sparse, particularly for long-range or inter-chromosomal interactions where interaction signal is weak. To mitigate the impact, we restrict borrowing to significant locus-locus interactions (interaction q-value *<* 0.05) and to loci located within ±1 Mb of the target locus. In addition, information borrowing is adaptively controlled by the data, including whether to borrow and how much to borrow. While these steps may not fully overcome limitations arising from Hi-C data quality, they provide a principled way to reduce its influence on the association test.

## Author Contributions

YH: Writing – original draft, Formal Analysis, Methodology, Conceptualization, Code implementation, Visualization; RB: Writing – original draft, review & editing, Methodology, Visualization; WL: Writing – review & editing, Conceptualization, Methodology; STH: Writing - review & editing, Code implementation; YL: Writing – review & editing, Conceptualization, Data curation, Methodology; JYT: Writing – review & editing, Conceptualization, Data curation, Methodology, Supervision, Project administration, Funding acquisition.

## Key Points

- Gene-based association tests leveraging 3D genome architecture have shown promise. Here we generalize this idea to gene-agnostic, whole-genome testing by introducing a Hi-C informed kernel association test, which allows a target variant set to borrow information from its Hi-C interacting loci when assessing its association with the trait.
- We present a principled procedure that converts Hi-C contact confidence into borrowing weights and embed these weights into genetic similarity kernels, so that higher-confidence interacting loci contribute more to the association test of the target variant set.
- We introduce a tuning parameter to govern how much the target locus borrows information from its interacting loci during association testing, so to ensure that only 3D interactions relevant to the trait under study are included.
- Using numerical analyses with real whole genome sequencing data, we show that integrating 3D genome architecture can improve association power and identify biologically meaningful variant associations that standard kernel association tests without Hi-C information may miss.

## Acknowledgments

**Atherosclerosis Risk in Communities (ARIC):** The Atherosclerosis Risk in Communities study has been funded in whole or in part with Federal funds from the National Heart, Lung, and Blood Institute, National Institute of Health, Department of Health and Human Services, under contract numbers (HHSN268201700001I, HHSN268201700002I, HHSN268201700003I, HHSN268201700004I, and HHSN268201700005I). The authors thank the staff and participants of the ARIC study for their important contributions.

**Genotypic/Genomic Dataset:** Molecular data for the Trans-Omics in Precision Medicine (TOPMed) program was supported by the National Heart, Lung and Blood Institute (NHLBI). Genome sequencing for “NHLBI TOPMed Whole Genome Sequencing (WGS) Project: ARIC” (phs001211.v5.p4) was performed at the Baylor College of Medicine Human Genome Sequencing Center (3U54HG003273-12S2, HHSN268201500015C). Core support including centralized genomic read mapping and genotype calling, along with variant quality metrics and filtering were provided by the TOPMed Informatics Research Center (3R01HL-117626-02S1; contract HHSN268201800002I). Core support including phenotype harmonization, data management, sample-identity QC, and general program coordination were provided by the TOPMed Data Coordinating Center (R01HL-120393; U01HL-120393; contract HHSN268201800001I). We gratefully acknowledge the studies and participants who provided biological samples and data for TOPMed.

**dbGaP Accession Number:** The datasets used for the analyses in this manuscript were obtained from dbGaP through dbGaP accession study numbers phs001211.v5.p4 and phs001211.v5.p4.c1.

## Data Availability Statement

The datasets analyzed for this study can be found in the NIH database of Genotypes and Phenotypes (dbGaP) through dbGaP accession study numbers phs001211.v5.p4 and phs001211.v5.p4.c1.

